# Chronic chromosome instability induced by Plk1 results in immune suppression in breast cancer

**DOI:** 10.1101/2022.06.16.496429

**Authors:** Sridhar Kandala, Lena Voith von Voithenberg, Sara Chocarro, Maria Ramos, Johanna Keding, Benedikt Brors, Charles D. Imbusch, Rocio Sotillo

**Affiliations:** Division of Molecular Thoracic Oncology, German Cancer Research Center (DKFZ), Im Neuenheimer Feld 280, 69120 Heidelberg, Germany; Faculty of Biosciences, Heidelberg University, Germany; Division of Applied Bioinformatics, German Cancer Research Center (DKFZ), Im Neuenheimer Feld 280, 69120 Heidelberg, Germany; Department of Medical Oncology, National Center for Tumor Diseases, Heidelberg University Hospital, Heidelberg, Germany; Translational Lung Research Center Heidelberg (TRLC), German Center for Lung Research (DZL)

## Abstract

Chromosomal instability (CIN), the inability to correctly segregate chromosomes during cell division, is a common characteristic of solid tumors. CIN contributes to tumor evolution by promoting intratumor heterogeneity, thus facilitating resistance to cancer therapies. *In vitro* studies have demonstrated that cells with complex karyotypes are recognized and eliminated by natural killer (NK) cells. Paradoxically, it has also been observed that human tumors with high levels of CIN have an immunosuppressive phenotype. It remains unclear which CIN-associated molecular features alter immune recognition during tumor evolution.

Previous studies with Polo-like kinase 1 (Plk1) overexpression in Her2-positive breast tumors, resulted in increased levels of CIN and delayed tumorigenesis. Using this mouse model, we show that high CIN tumors activate a senescence-associated secretory phenotype (SASP) and become immune evasive by activating RELB signaling and upregulating PD-L1 in a non-cell-autonomous manner. Single-cell RNA sequencing of immune cells from early-stage induced mammary glands revealed that macrophages, NK cells, B cells and regulatory T cells are programmed to a suppressive phenotype during tumor development. In human tumors, we further establish the importance of RELB/p38 signaling in understanding the interplay between CIN and the immune system, highlighting the need for novel adjuvant therapies in the context of chromosomally unstable tumors.

## Introduction

Frequent chromosome missegregation is a hallmark of human cancers (Hanahan, 2022). Several mechanisms such as defects in cell-cycle regulation or alteration of the function of the spindle assembly checkpoint (SAC), compromise mitotic segregation fidelity to induce chromosome instability (CIN) (Jaarsveld & Kops, 2016; Potapova et al., 2013). To date, several mitotic proteins have been reported to be upregulated in different tumor types, including PLK1, a member of the Polo-like kinase family. We have previously shown that overexpression of Plk1 in a Her2-driven breast cancer mouse model results in polyploidy, increased levels of CIN, and a significant delay in breast tumor formation with a good prognosis for cancer patients (Carcer et al., 2018).

At the same time, CIN can also drive metastasis, confers drug resistance and is associated with poor patient prognosis in multiple cancer types (Lukow et al., 2021). To explore CIN as a therapeutic target and to perceive its role in disease progression, it is critical to understand how different levels of CIN affect tumor development and its microenvironment, further contributing to therapy resistance. Analysis of TCGA data across 12 different human cancer types revealed that high aneuploid tumors show reduced expression of markers for cytotoxic cells (CTLs) and Natural Killer cells (NK cells) and increased infiltration of tumor-promoting macrophages (M2 TAMs), resulting in decreased antitumor immunity (Davoli et al., 2017). Moreover, recent discoveries show that multiple immune effectors along with interleukin/toll-like receptors mediated by the innate immune system participate in sensing hyperploid cells (Aranda et al., 2018; Senovilla et al., 2012). Similarly, cells with complex karyotypes express high levels of the NK cell ligand *NKG2D* such as *MICA, ULBPs* that mediate NK cell activation (Santaguida et al., 2017) or induce micronuclei formation and activate the cGAS-STING pathway facilitating their clearance (Dou et al., 2017; Vizioli et al., 2020).

In human tumors, it has been shown that PLK1 expression induces inflammation and correlates positively with the number of tumor-infiltrating lymphocytes (TILs) (M. Li et al., 2018; Zhou et al., 2021). However, the interplay between aneuploid tumor cells and the tumor microenvironment in the context of CIN induced by PLK1 still remains obscure. Here, we investigate the consequences of CIN induced by Plk1 overexpression *in vivo* in a breast cancer mouse model. Plk1 overexpression triggered the formation of a senescence-associated secretory phenotype (SASP) ultimately resulting in immune evasion of the tumors through upregulation of PD-L1 in a non-cell-autonomous manner mediated by RELB and p38a. Interestingly, single-cell RNA sequencing of early-stage tumors identified the presence of Arg1^+^ TAMs, decrease in cytotoxicity of NK cells and increased infiltration of regulatory T cells (Tregs) in high CIN tumors. These may contribute to a suppressive phenotype and thus lead to diminished antitumor responses. Our results also show that human breast tumors harboring high levels of *PLK1* display an immune-suppressive phenotype associated with T-cell exhaustion, and SASP driven by NF-kB/ p38 signaling, suggesting that modulation of NF-kB activity or immune checkpoint inhibition could be used as adjuvant therapy in inflammatory tumors expressing high levels of *PLK1*.

## Results

### High CIN results in a Senescence Associated Secretory Phenotype

CIN is one of the key mechanisms inducing immune evasion in tumors (Bakhoum et al., 2018; Burrell et al., 2013; Davoli et al., 2017; McGranahan et al., 2017; Rosenthal et al., 2019; Watkins et al., 2020). How different levels of CIN shape and influence tumorigenesis in the context of an active immune system is unclear. To uncover this, we used a breast cancer mouse model that overexpresses Her2 (Her2) or Her2 and Plk1 (Her2-Plk1) by using the reverse tetracycline transactivator under the control of the mouse mammary tumor virus promoter (MMTV-rtTA) (Carcer et al., 2018). We performed low-pass, whole-genome sequencing on the resulting primary breast tumors (median coverage 0.11-fold), followed by somatic copy number alteration (SCNA) analysis. Her2-Plk1 tumors had elevated levels of SCNA (high aneuploidy) compared to Her2 tumors (low aneuploidy) [Fig1A-C, FigS1A]. As elevated levels of Plk1 are expected to drive a CIN phenotype, we performed time-lapse microscopy of Her2 and Her2-Plk1 breast tumor cells. Her2 tumor cells display 24% of mitotic errors compared to 50% in Her2-Plk1 cells [FigS1B]. Moreover, Plk1-overexpressing tumor cells displayed a higher frequency of aneuploidy and polyploidy compared to Her2 tumor cells (Carcer et al., 2018). Altogether these results suggest that in mammary tumors Plk1 overexpression results in increased CIN.

**Figure 1:**
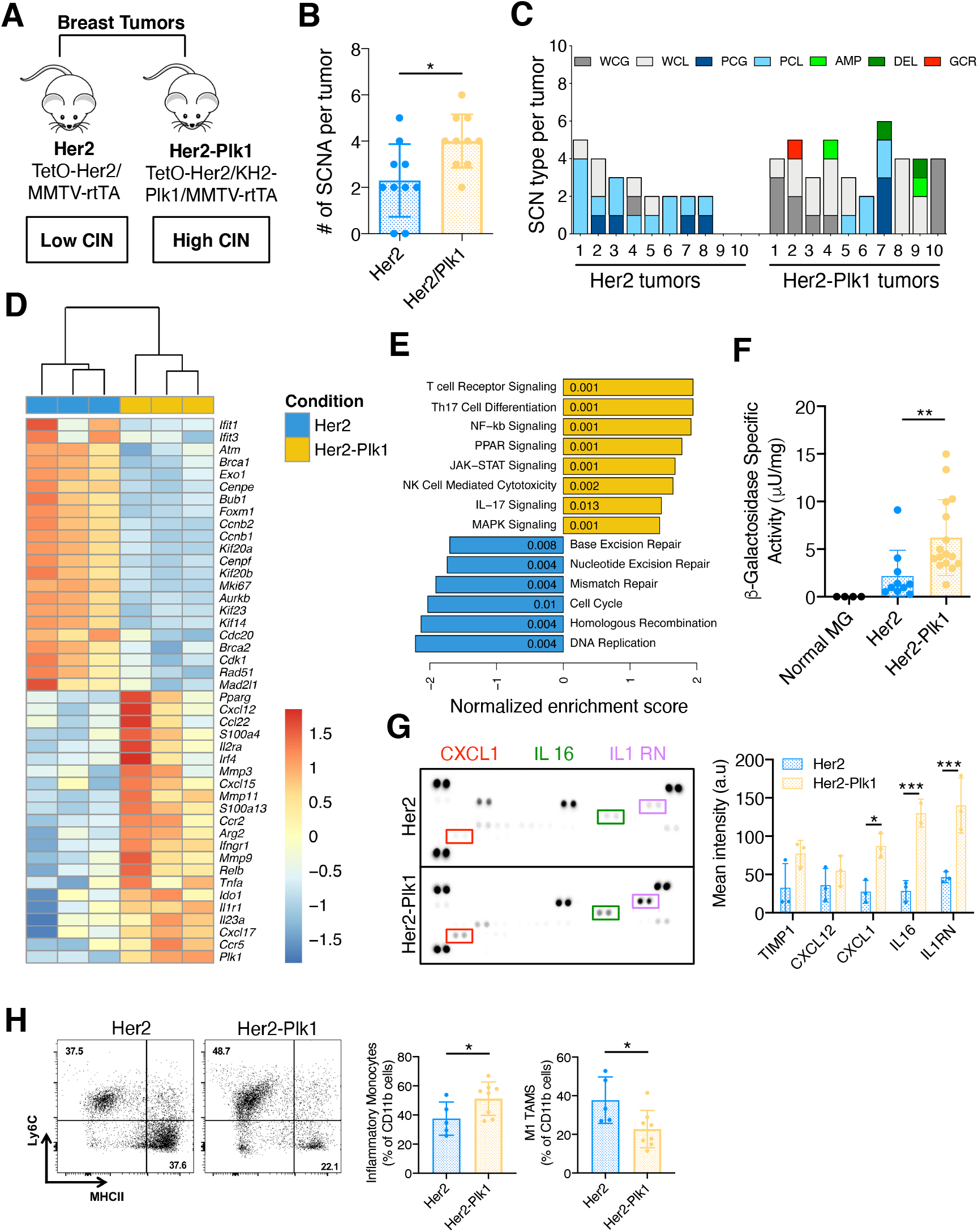
Plk1 overexpression drives inflammatory monocyte recruitment as a consequence of SASP. **A)** Schematic representation showing the mouse models used in this study, where Her2 mammary tumors show low levels of CIN and Her2-Plk1 overexpression results in high CIN tumors. **B**) Average SCNAs in Her2 and Her2/Plk1 tumors; Unpaired t test, p = 0.012. **C**) SCNAs in 10 Her2 and 10 Her2/Plk1 mammary tumors. Shown are whole chromosome gain (WCG) and loss (WCL), partial chromosome gain (PCG) and loss (PCL), focal amplification (AMP), deletion (DEL) and gross chromosomal rearrangement (GCR). **D)** Total transcriptome of Her2 and Her2-Plk1 tumor tissues was analyzed by bulk RNA-sequencing. Heatmap showing differentially expressed cell cycle and SASP-related genes (Fold change (FC) > 1.5 or < −1.5, *p*<0.05, n=3 tumors per genotype). **E)** GSEA using gene sets from KEGG pathways as analyzed by cluster profiler. Pathways overrepresented in Her2-Plk1 tumors are shown in yellow, whereas pathways overrepresented in Her2 tumors are displayed in blue. **F)** Fluorometric measurement of β-galactosidase activity in Her2 and Her2-Plk1 tumor lysates (Mann Whitney test p<0.005**, n=10 (Her2) and n=16 (Her2-Plk1)). **G)** Cytokine array of tumor lysates of Her2 and Her2-Plk1 showing increased CXCL1, IL16 and IL1RN in the high CIN group (p<0.05* p<0.0005***, 2-way ANOVA, Sidak’s multiple comparison test, n=3). **H)** Characterization of TAM’s and monocytes. Live CD45^+^CD11b^+^ cells were divided into two groups based on Ly6C and MHCII. Flow cytometric analysis of inflammatory monocytes (Ly6c^high^MHCII^low^) and M1 TAMs (Ly6c^low^MHCII^high)^ in endpoint tumors (p<0.05*, n=5 (Her2) and n=8 (Her2-Plk1)).

High CIN levels in primary mammalian cells have been reported to display senescence-associated features and upregulation of inflammatory cytokines (Andriani et al., 2016). Therefore, to explore the immune signatures associated with Plk1 overexpression, we performed whole transcriptome analysis by RNA-sequencing of these Her2 and Her2-Plk1 breast tumors at a humane endpoint. Hierarchical clustering was employed to understand the similarity between various samples across both groups. The results of this clustering determined that Her2-Plk1 segregate away from Her2 tumors [FigS1C]. Differential expression analysis revealed that Plk1 overexpressing tumors had higher expression of several SASP markers, characterized by the upregulation of cytokines/chemokines (*Tnfa, Ido1, Cxcl15, Cxcl17, Cxcl12*) and extracellular matrix ECM proteases (*Mmp3, Mmp9, Mmp11*), and the downregulation of genes associated with cell cycle regulation (*Bub1, Kif20a, Kif20b, Cenpe, Aurkb, Mad2l1*) and DNA damage repair (*Brca1, Brca2, Atm, Exo1, Rad51*) [Fig1D]. Gene set enrichment analysis (GSEA) using gene sets from Kyoto Encyclopedia of Genes and Genomes (KEGG) pathways and Gene Ontology (GO) analysis, identified significant enrichment of signatures related to inflammation (Il17 signaling, JAK-STAT signaling) and SASP (NF-kB signaling), and downregulation of DNA damage response signatures (base excision repair, nucleotide excision repair, mismatch repair, homologous recombination) in the high CIN group [Fig1E, FigS1D].

We next asked whether Plk1 overexpression results in the senescence of tumor cells. Lysates from Her2 and Her2-Plk1 tumors showed increased ß-galactosidase activity in Her2-Plk1 tumors compared to Her2, confirming the increase in senescence in the high CIN group [Fig1F]. These findings correlate with our previous observation in mouse embryonic fibroblasts, where Plk1 overexpression resulted in defective cell proliferation, formation of polyploid cells and ultimately senescence (Carcer et al., 2018). Additionally, we performed a cytokine array of Her2 and Her2-Plk1 tumors and found an increased expression of SASP related inflammatory cytokines such as CXCL1, IL16 and Il1RN in the latter group [Fig1G], further supporting our previous observations. As inflammation is known to recruit monocytes that circulate through the blood and extravasate (Shi & Pamer, 2011), we performed flow cytometry to investigate monocyte influx in response to SASP. Inflammatory monocytes and M1 macrophages were gated from CD11b^+^ cells using a previously described strategy (Georgoudaki et al., 2016). We detected a significant increase in CD11b^+^Ly6C^high^MHCII^low^ inflammatory monocytes and a reduction in CD11b^+^Ly6c^low^MHCII^high^ M1 tumor-associated macrophages (TAMs) in Her2-Plk1 tumors compared to Her2 tumors [Fig1H]. Taken together, these results suggest that CIN induced by Plk1 overexpression leads to an influx of inflammatory monocytes as a consequence of SASP.

### High CIN activates non-canonical NF-kB (RelB) signaling and promotes immune suppression in a non-cell-autonomous manner

Chromosome missegregation leads to chronic stress, endogenous DNA damage and activation of senescence (Q. He et al., 2018; Sheltzer et al., 2016; Torres et al., 2007). NF-kB being a master transcription factor of senescence and inflammation drives immune sensing of chromosomally unstable cells through the recruitment of NK cells (Wang et al., 2021) and also mediates immune evasion by promoting migration, invasion and metastasis through non-canonical RELB signaling (Bakhoum & Cantley, 2018). Furthermore, NF-kB regulates PD-L1 expression through the release of inflammatory cytokines like TNFα by TAMs or IFNγ by infiltrating T and NK cells (Antonangeli et al., 2020; Budczies et al., 2017). We, therefore, asked whether Plk1 overexpression could activate NF-kB dependent PD-L1 expression as a consequence of inflammation. To answer this question, we analyzed the protein levels of PLK1, NF-kB-p65, RELB, p38a and phospho-p38 in Her2 and Her2-Plk1 breast tumors samples. We found significantly higher protein levels of all these proteins (indicative of non-canonical signaling) in the high CIN group [Fig2A]. Importantly, MCF7 and Cal51-rtTA cells, in isolation without a surrounding microenvironment, infected with a doxycycline-inducible PLK1 vector did not show any difference in NF-kB expression levels after induction, suggesting the importance of immune cells in the tumor microenvironment in mediating this effect [FigS2A and B]. To confirm that the increased expression of NF-kB was not merely due to Plk1 expression, we used another breast cancer model that overexpresses Mad2 (Rowald et al., 2016)Protein lysates from Her2-Mad2 tumors showed a similar increase of NF-kB-p-65, RELB and phospho-p38 [FigS2C], suggesting that this activation was not due to elevated levels of Plk1 but rather a direct consequence of CIN.

**Figure 2:**
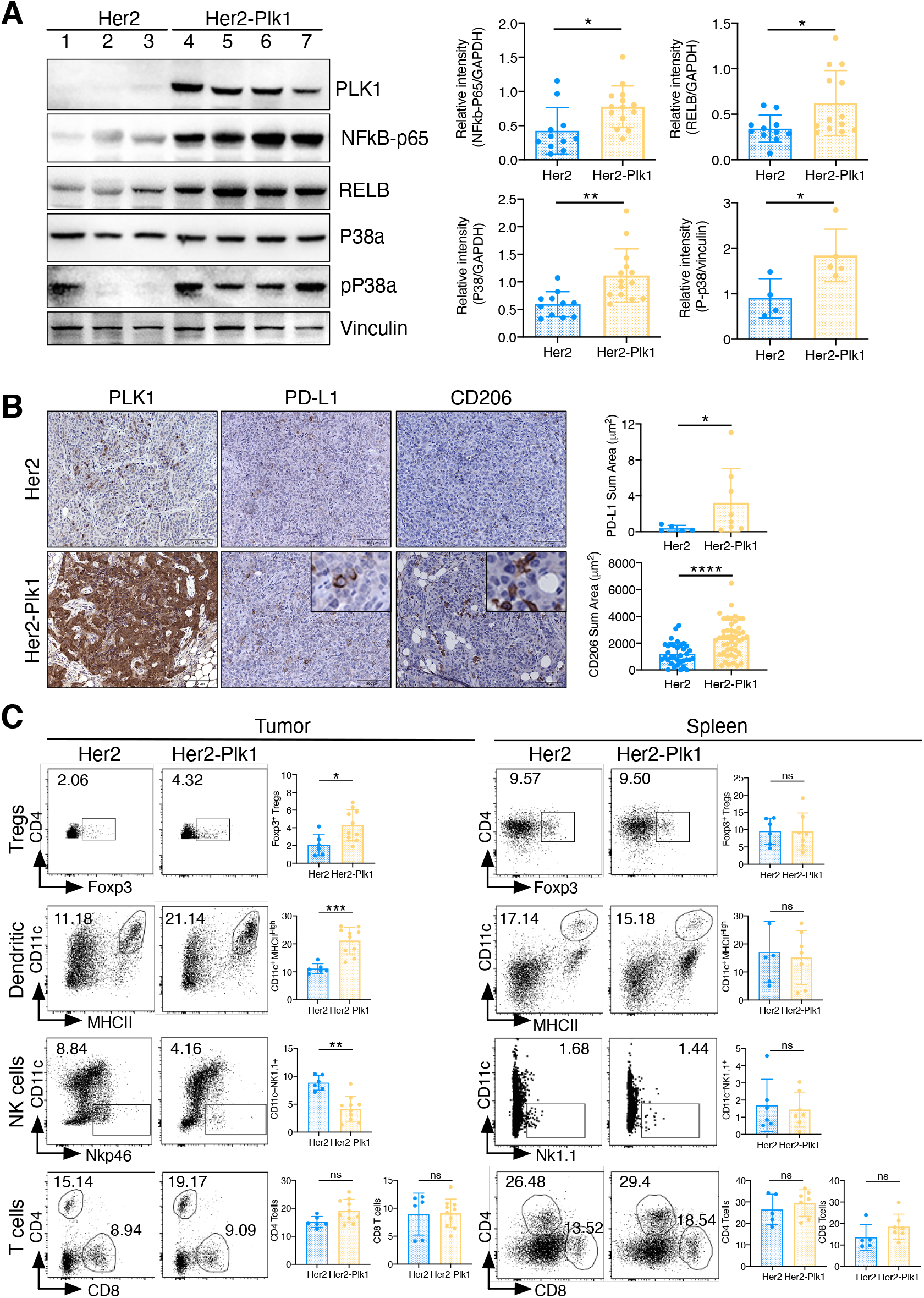
Plk1 overexpression mediates immune suppression as a consequence of non-cell-autonomous effects. **A)** Expression of P38a and NF-kB-p65 in breast tumors from Her2 and Her2-Plk1 mice. Tumor cell lysates were used for immunodetection by using anti-PLK1, anti-NFkB-p65, anti-RELB, anti-P38, anti-pP38 and anti-vinculin as a loading control. Bar graphs represent the relative intensity of the signal from immunodetection (*, p<0.05, **, p<0.005, Mann-Whitney test, n=11 (Her2), n=14 (Her2-Plk1); numbers represent different tumors). **B)** Immunohistochemistry of PLK1, PD-L1 and CD206 in Her2 and Her2-Plk1 tumor sections and their corresponding bar graphs representing total sum area (μm^2^). Positive regions per sample was calculated by dividing DAB positive area in each ROI/total area of the ROI (region of interest) and total sum area (μm^2^) and adding positive regions in all the individual samples. Scale bar: 100μm. (*, p<0.05, **** p< 0.0001, Mann-Whitney test, n=5 (Her2), n=8 (Her2-Plk1)). **C)** Quantification of tumor-infiltrating lymphocytes in Her2 and Her2-Plk1 tumors and spleens. Representative two-parameter dot plots of CD45^+^ gated Tregs (CD25^+^CD4^+^Foxp3^+^), dendritic cells (CD11c^+^MHCII^high^), NK cells (CD11c^−^Nkp46/Nk1.1^+^)) and T cells (CD3^+^CD4^+^, CD3^+^CD8^+^) in both spleen (n=5 (Her2), n=7 (Her2-Plk1)) and tumors (*, p<0.05, ** p<0.005, ***, p< 0.0005, Mann-Whitney test n=6 (Her2), n=10 (Her2-Plk1)) and bar graphs represented as % of CD45 positive cells. The numbers depicted in the boxes refer to the percentage of each cell type in different samples (Her2 and Her2-Plk1).

Cell-autonomous effects associated with overexpression of Plk1 induced tumor-suppressive effects in a breast tumor model *in vivo* and the activation of cell cycle arrest in G1 phase mediated by p21 (Carcer et al., 2018). To address how senescence and SASP mediated non-cell-autonomous effects alter the tumor microenvironment in high CIN tumors, we performed immunohistochemistry for PD-L1 and CD206, an (M2 macrophage marker), in mouse breast tumor sections [Fig2B]. Her2-Plk1 tumors displayed elevated levels of PD-L1 and CD206, suggesting a suppressive phenotype in these tumors. More complete immunophenotyping of breast tumor tissues [FigS2D] as well as from spleens revealed an increased infiltration of Tregs (CD3^+^CD4^+^CD25^+^Foxp3^+^) and dendritic cells (CD11c^+^MHCII^high^) and a decrease in the total number of infiltrating NK cells (CD11c^−^Nkp46^+^/Nk1.1^+^) in the Her2-Plk1 breast tumors [Fig2C], which was not observed in the spleen of these mice. These results indicate that the recruitment of Tregs and reduced infiltration of NK cells is a direct result of Plk1 overexpression resulting in higher CIN. Altogether, these data suggest that high CIN upregulates non-canonical NF-kB signaling and suppresses antitumor immunity as a key consequence of non-cell-autonomous effects.

### Immune modulation of innate cells occurs at an early stage during the development of high CIN tumors

To understand how CIN shapes the immune microenvironment to a suppressive phenotype, we collected mammary glands of Her2 and Her2-Plk1 mice at early stages of tumor development, in a pre-malignant stage, which we found to be after 16 days on doxycycline in Her2 mice and 22 days in Her2-Plk1 [Fig3A, FigS3A]. We first performed PLK1 staining and fluorescence *in situ* hybridization of these mammary glands to determine the aneuploidy levels in both groups. Similar to humane endpoint tumors (Carcer et al., 2018) and Fig1B and Fig1C, the Her2-Plk1 mammary glands at early stages showed an increase in PLK1 expression and aneuploid levels compared to Her2 mammary glands [FigS3B]. To assess the immunosuppressive phenotype, induced by CIN, we performed single-cell mRNA sequencing (scRNA-Seq) on 23120 CD45^+^ hematopoietic cells [TableS1, FigS3C] sorted from these early doxycycline-induced mammary glands of Her2 and Her2-Plk1 mice. We analyzed the data using the Seurat package (Butler et al., 2018) and performed cell-type annotation based on SingleR and the ImmGen database (Consortium et al., 2008) to identify the different innate and adaptive populations [Fig3B] and determine their relative number [FigS3D and FigS3E]. The tumor microenvironment of all samples was dominated by T cells (approx. 60-70% of all cells). A comparison between the subsets with high and low CIN revealed that the relative number of T cells (especially CD4 T cells and Tregs) was higher in the Her2-Plk1 group, whereas the relative number of macrophages and NK cells was lower. *In vitro* experiments have shown that aneuploidy also affects the polarization of macrophages impairing antigen-specific T cell activation (Xian et al., 2021). Since CIN induces inflammation and facilitates tumor progression by recruiting macrophages and neutrophils (Duijf & Benezra, 2013; M & E, 2019; Roschke & Rozenblum, 2013), we analyzed the macrophage population in the Her2 and Her2-Plk1 samples and identified two subsets of TAMs. While the first subset was characterized by CD11b^+^CD11c^−^ macrophages overexpressing *Cd209d, Retnla/Fizz1, Mrc1, F13a1, Clec10a, Ccl8, Selenop*, and *Igfbp4*, the second subset consisted of CD11b^+^CD24^−^ macrophages overexpressing *Fn1, Ctss, Hexb, Gatm, Ctsl*, and *Runx3* [FigS4A]. *Tnf* superfamily genes such as *Tnf, Tnfrsf1a, Tnfrsf1b* and genes activated in response to *Tnf* mediated inflammation such as *Il1b, Nlrp3, Nfkb1, Traf2, Tlr2, Tmem173* were expressed in both subsets of early tumor groups irrespective of their ploidy status [FigS4B]. Differential gene expression of the CD11b^+^CD11c^−^ macrophage subset showed decreased expression of genes involved in the maintenance of proinflammatory transcriptional signaling, such as *S100a8/S100a9, Tpt1, Ltb, Crip1, Smad7, Ccl24, Ccl6* and *Ccr7* in Her2-Plk1 [Fig3C]. Similarly, CD11b^+^CD24^−^ macrophages from Her2-Plk1 mammary glands presented an upregulation of *Arg1, Gpnmb, Il1rn, Fcgr2b, Pltp, Pdpn, Adam8, Ahnak, Fam20c, Nrp2 and Timp1* [Fig3D], all of which have been associated with wound healing, epithelial to mesenchymal transition (EMT) and a migratory phenotype. Additionally, they showed reduced expression of the MHCII-related genes *H2-ab1, H2-Aa, H2-Dma, H2-eb1, H2-Dmb2, Ciita* and *Cd74* corresponding to decreased antigen presentation [Fig3D]. Altogether, these results suggest that CIN induced by Plk1 expression leads to an early increase in *Arg1*^*+*^ TAMs thereby dampening macrophage effector functions by decreasing proinflammatory cytokine production and antigen presentation.

**Figure 3:**
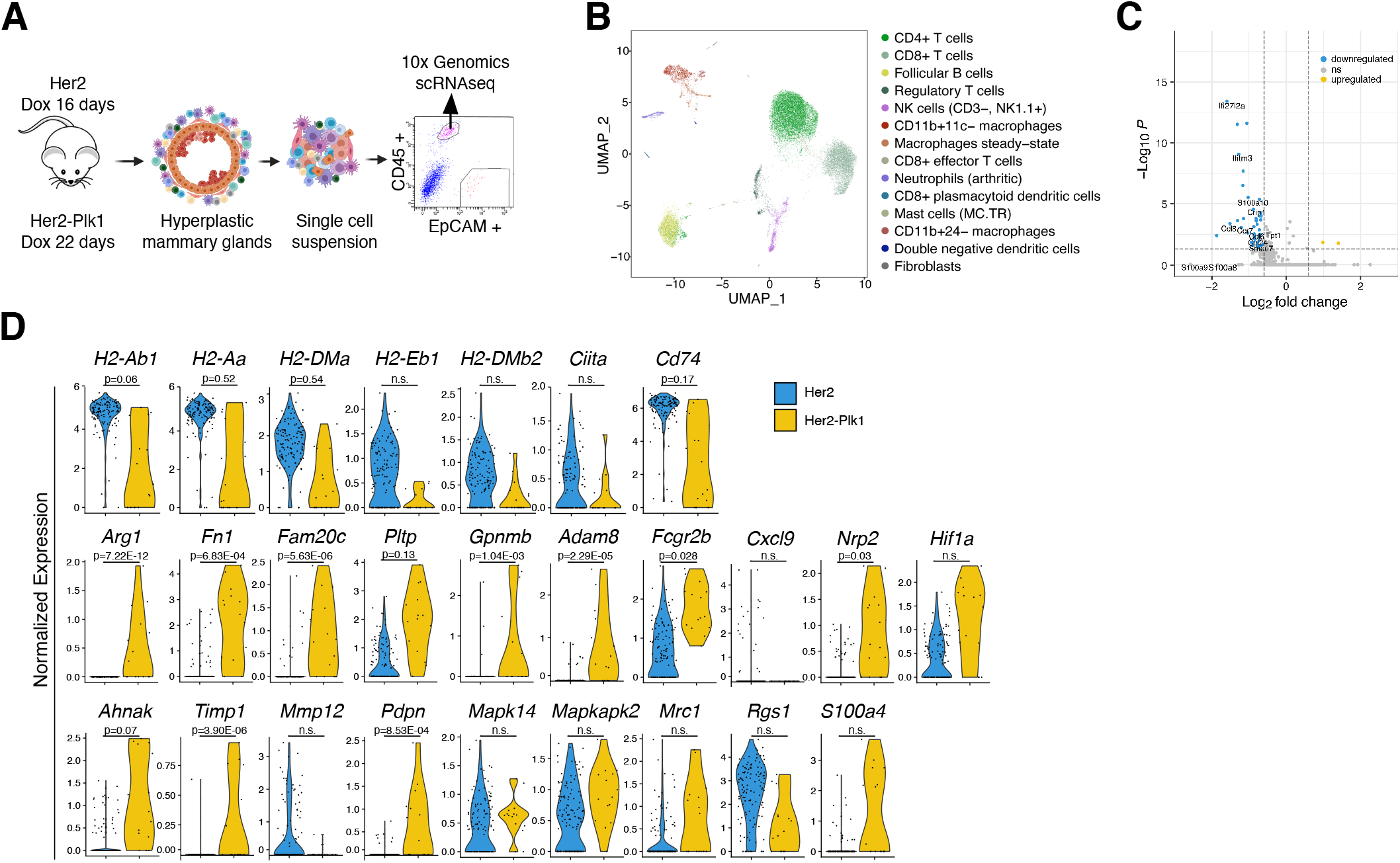
Innate immune cell phenotype of high and low CIN tumors as investigated by single-cell RNA-sequencing. **A)** Schematic depiction of isolation of hyperplastic mammary glands and FACS sorting of individual cells to obtain CD45 hematopoietic cells at early stage Her2 (16 days on doxycycline) and Her2-Plk1 (22 days on doxycycline) samples. Samples were extracted from three different mice per genotype from two mammary glands each. **B)** Clustered immune cell populations displayed by UMAP in all samples. The cell types in the clusters were annotated by using SingleR and gene expression information from the ImmGen reference database (Consortium et al., 2008) and displayed in a color-coded fashion. **C)** Volcano graph showing differentially expressed genes in the CD11b^+^CD11c^−^ macrophage subtype with log2-fold change in gene expression versus -log10 of adjusted p-values. Genes that were found to be downregulated in Her2-Plk1 with respect to Her2 are shown in blue, while genes that were upregulated are shown in yellow (by a fold change of at least ±0.6 and an adjusted p-value of ≤0.05). **D)** Violin graphs showing the normalized expression levels of CD11b^+^CD24^−^ macrophages for genes related to antigen presentation, EMT and anti-inflammatory wound-healing phenotypes. Her2 samples are in blue and Her2-Plk1 in yellow. p-values adjusted for multiple testing are shown above each graph. Each dot represents the gene expression level of an individual cell.

NK cells play a prominent role in the detection of cells with aberrant chromosome numbers and have been shown to eliminate them *in vitro* (Santaguida et al., 2017; Wang et al., 2021). To functionally validate the role of NK cells in immune sensing of aneuploid tumor cells *in vivo*, we treated Her2 and Her2-Plk1 animals with an NK blocking antibody until the appearance of a palpable tumor. We observed that although Her2 tumors grew faster upon depletion of NK cells this difference was not significant. However, Her2-Plk1 animals treated with the NK block presented a significant reduction in survival time [Fig4A].

**Figure 4:**
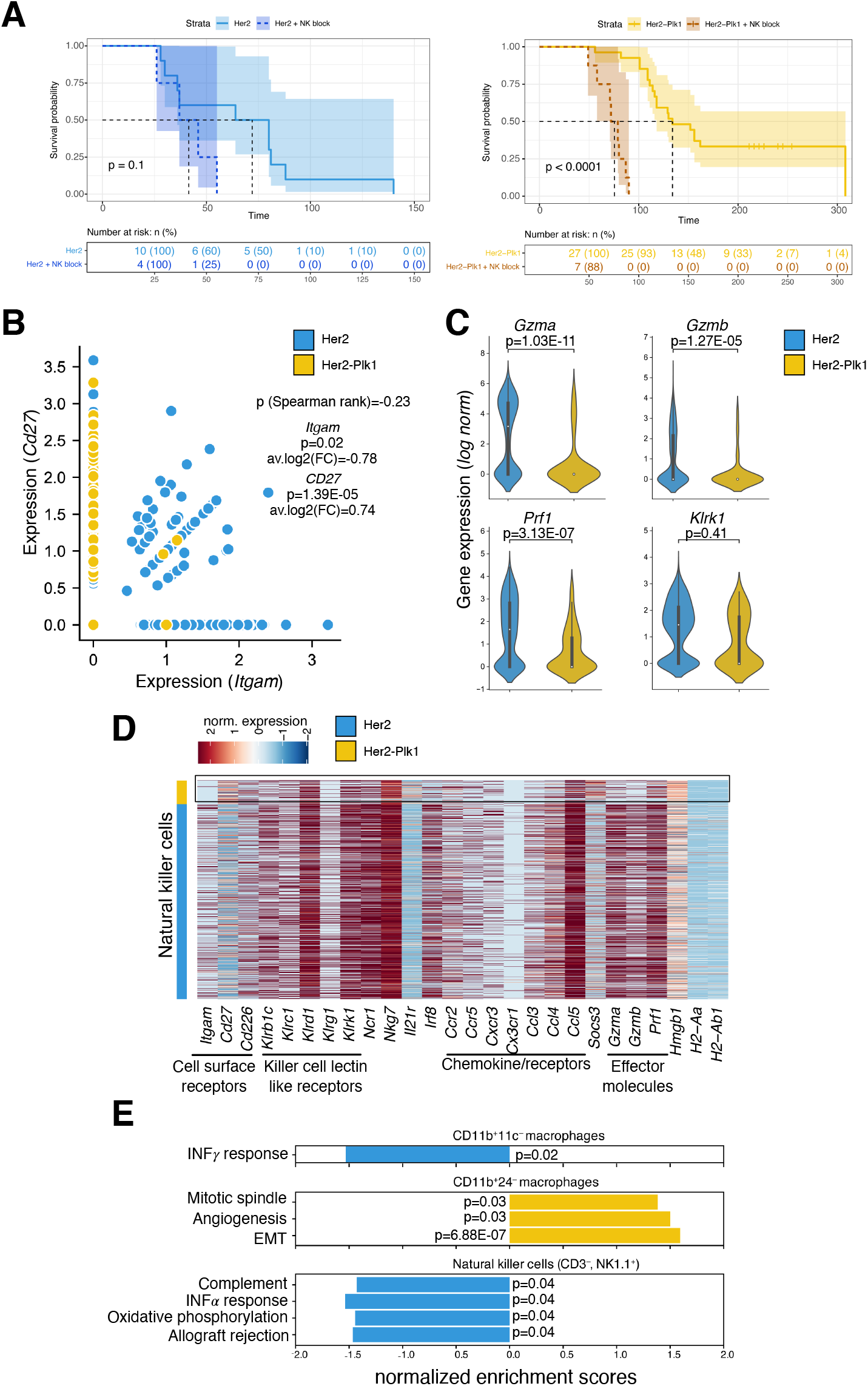
Role of NK cells in immune response of high CIN tumors. **A**) Survival analysis of Her2 (left) and Her2-Plk1 (right) tumors of mice treated with NK1.1 blocking antibody. Survival probability over time (upper graphs) and number and percentage of mice alive over time (lower table). The curves in light blue and light yellow depict untreated mice, whereas the curves in dark blue and dark yellow show mice treated with NK block, respectively. **B)** Normalized gene expression of *CD27* versus *Itgam* (*CD11b*) in NK cells at the single cell level (individual dots) (Her2: blue, Her2-Plk1: yellow). The Spearman rank correlation coefficient across samples as well as the fold change and p-values adjusted for multiple testing for differential expression between Her2 and Her2-Plk1 samples are provided. **C)** Violin graphs showing the log-normalized gene expression values of NK cells in Her2 (blue) and Her2-Plk1 (yellow) tumors exemplarily shown for genes involved in cytotoxic response and surface receptors of NK cells. **D)** Heatmap showing normalized gene expression levels of differentially expressed effector genes, chemokines and their receptors, and activating receptor genes in NK cells. Each line represents a single cell. **E)** Gene set enrichment analysis for identification of pathways with enrichment of differentially expressed genes for two macrophage subsets and NK cells. Y-axis: hallmark signatures and X-axis: normalized enrichment scores with adjusted p-values, color-coded by the direction of the effect (yellow: upregulation in Her2-Plk1, blue: upregulation in Her2).

From the single-cell analysis, we observed differential expression of *Cd27* and *Itgam* (*CD11b*) between NK cells from Her2 and Her2-Plk1 samples [Fig4B]. This functional classification suggests that the NK cells in Her2-Plk1 tumors show a lower cytotoxicity (Abel et al., 2018). Interestingly, we additionally observed a similar trend with downregulation of NK-effector molecules like *Gzma, Gzmb* and *Prf1*, whereas the expression of activating receptor *Klrk1* (NKG2D) was not significantly different between both groups [Fig4C]. Furthermore, compared to Her2-Plk1, Her2 samples showed an upregulation of other NK-specific effector molecules such as *Nkg7*, chemokines (*Ccl3, Ccl4, Ccl5)* and activating receptors (*Klrb1c, Ncr1, Klrd1, Cd226)* compared to Her2-Plk1 samples [Fig4D]. The development, functional maturation and trafficking of NK cells in samples with high CIN were affected by *Irf8, Socs3, Cx3cr1* and hypoxia [Fig4D]. Gene set enrichment analysis of differentially expressed genes between NK cells in Her2 versus Her2-Plk1 samples revealed the downregulation of hallmark signatures of IFNα response, oxidative phosphorylation, and the complement system in high CIN samples [Fig4E]. Similarly, GSEA of CD11b^+^24^−^ macrophages showed pathways related to the mitotic spindle, angiogenesis and EMT upregulated in Her2-Plk1 samples [Fig4E]. Thus, our data suggest early modulation of macrophages to Arg1^+^ TAMs and limiting effector NK cells thereby contributing to diminished anti-tumor responses against tumors with high levels of CIN.

### Plk1 high tumors show an autoimmune phenotype mediated by B cells and an increased infiltration of resting Tregs

B cells are an important component of the adaptive immune system that mediate humoral immunity. B cells inhibit tumor development through tumor-reactive antibodies and aid in NK cell killing, macrophage phagocytosis and CD8 T-cell cytotoxicity. However, they can contribute to tumor growth through the production of autoantibodies (Lee et al., 2021) and differentiation into regulatory B cells (Y. He et al., 2014). To illustrate the characteristics of B cells in our model, we performed differential expression analysis of the B cell cluster between Her2 and Her2-Plk1 samples. B cells from high CIN tumors showed a decreased expression of *Mzb1, Ltb, C1qa, Pglyrp1, Eif3d, Ccl6, Myl12b, Ccnd3, Ighg1, Cd47, Ptpn6, Rgs1, S100a10, Gimap1, Cd3g, Irf8, Ptpn22, Scd1* and *Cd79b* that affect the B cell development, activate autoreactive B cells and aid in the production of anti-DNA antibodies thereby contributing to an autoimmune phenotype [Fig5A]. Studies have shown that *Cd79, Mzb1, Scd1* and *Ptpn22* are associated with dysregulated downstream signaling of B cell receptor (BCR), autoantibody production and altered lipid metabolism (Hardy et al., 2014; Miyagawa-Hayashino et al., 2018; Zhang et al., 2021). The phenotype presented here suggests an essential role of B cells in triggering autoimmune responses in tumors with high Plk1, although the functional validation of an autoimmune phenotype in this mouse model remains to be elucidated.

**Figure 5:**
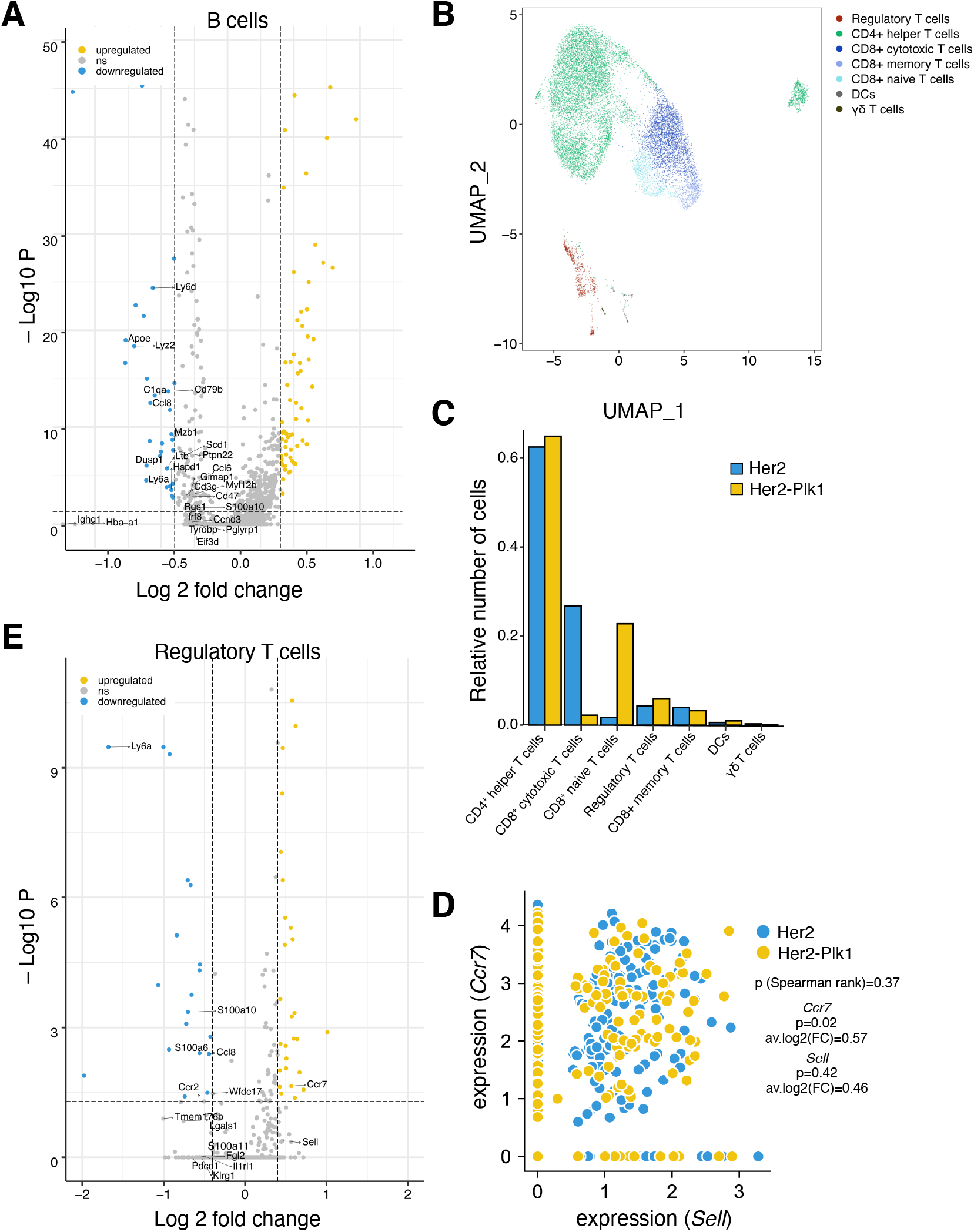
B- and Treg-mediated adaptive immune response in high CIN tumors. **A)** Volcano graph with reduced axes to focus on selected differentially expressed genes (labeled) in the B cell subset of Her2 and Her2-Plk1 tumors displayed as log2 fold change versus - log10 of the adjusted p-value. Genes that were found to be downregulated in Her2-Plk1 with respect to Her2 are shown in blue, while genes that were overexpressed are shown in yellow (by a fold change of at least ±0.4 and an adjusted p-value of ≤0.05). **B)** UMAP graph of T cells clustered into different T cell subtypes. Seven clusters with different cell types were identified after cell type annotation. **C**) Relative distribution of the number of cells in each of the T cell subtypes with Her2 in blue and Her2-Plk1 in yellow. **D)** Normalized gene expression of *Sell* (*Cd62L*) versus *Ccr7* in regulatory T cells at the single cell level (individual dots) (Her2: blue, Her2-Plk1: yellow). The Spearman rank correlation coefficient across samples as well as the fold change and p-values adjusted for multiple testing for differential expression between Her2 and Her2-Plk1 samples are provided. **E)** Volcano graph with reduced axes to focus on selected differentially expressed genes in the cluster of regulatory T cells displayed as log2 fold change versus -log10 of the adjusted p-value. Genes downregulated in Her2-Plk1 compared to Her2 are shown in blue, genes upregulated in Her2-Plk1 are displayed in yellow (by a fold change of at least ±0.4 and an adjusted p-value of ≤0.05).

To further characterize T cell infiltration in high CIN samples, we performed sub-clustering of T cells in Her2 and Her2-Plk1 samples. We identified five distinct T cell phenotypes (Treg cells, CD4^+^ helper T cells, CD8^+^ cytotoxic T cells, CD8^+^ memory T cells, and CD8^+^ naïve T cells) and two additional sub clusters with no conclusive information due to their low number of cells [Fig5B] and their relative proportions [Fig5C]. In the Her2-Plk1 samples, we observed an increase in the relative number of CD8^+^ naïve T cells, regulatory T cells, and CD4^+^ helper T cells in comparison to the total T cell number, whereas a decrease in the number of CD8^+^ cytotoxic T cells was seen. Under conditions of chronic inflammation, Tregs are recruited to the inflamed tissue as an extrinsic mechanism to keep immune homeostasis by maintaining self-tolerance and inhibiting excessive immune responses to self-antigens. Regulatory T cells can modulate the immune system and suppress the induction and proliferation of effector T cells, thereby dampening antitumor immunity (Mougiakakos et al., 2010). Differential expression analysis of regulatory T cells revealed a slight upregulation of the expression levels of *Sell* (CD62L) (FC=0.5, p_adj_=0.4) and *Ccr7* (FC=0.6, p_adj_=0.02) [Fig5D] in Her2-Plk1 tumors compared to Her2, suggesting a decrease of the effector Treg phenotype and an increase in the resting phenotype. In regulatory T cells of Her2-Plk1 tumors, we observed a downregulation of the genes *Ccr2, Ccl5*, and *Ccl8*, affecting Treg function, compared to Her2 tumors [Fig5E]. It is known that a higher expression of *Ccr2* in Tregs correlates with increased activity of these cells (Zhan et al., 2020). Thus, our data suggest a decrease in Treg activity and an increase in the relative number of resting Tregs in high CIN tumors.

### Human breast tumors expressing high levels of PLK1 are predicted to respond to adjuvant therapies involving NF-kB or PD-L1 inhibition in combination with first-line treatments

To expand our analysis of the relationship between PLK1 expression and immune cell composition in different types of human breast tumors, we obtained gene expression and copy number data from 1098 TCGA breast tumors. These were further classified as PLK1-high (Z-score >1) and PLK1-low (Z-score < 1) based on the PLK1 expression (log2 (normalized counts +1)). The average expression of PLK1 in the cohort was 8.817 (SD of 1.32) [Fig6A]. Consistent with our previous *in vivo* findings, we saw PLK1-high cohort harboring higher levels of chromosome gains and losses [FigS5A]. Unsupervised clustering using principal component analysis was done to understand the variance between 186 PLK1-high and 165 PLK1-low tumors [Fig6B]. Higher expression of PLK1 correlated with aggressive basal-like tumors (171 basal tumors out of 186), whereas lower levels of PLK1 were associated with luminal breast cancers (84 luminal tumors out of 165) [Table S2]. This result is in line with previous studies where PLK1 inhibition was shown to be a potential target for the treatment of triple negative breast cancers (Giordano et al., 2019; Nieto-Jimenez et al., 2020; Ueda et al., 2019). We then used the cell deconvolution tool CIBERSORT to estimate the relative number of tumor-infiltrating lymphocytes, macrophages, dendritic cells and NK cells from PLK1 low and high groups using the gene expression data. Following our data for our mouse model, PLK1 high tumors showed increased percentages of Tregs, M0/M1 macrophages and reduced infiltration of naïve B-cells and CD8^+^ /CD4^+^ T cells resulting in immune exclusion [Fig6C]. Moreover, expression data of genes corresponding to signatures of immune suppression (*ARG2, CD274, IL10, IDO1, PDCD1, PDCD1LG2, VEGFA, VEGFB, CTLA4 and TGFB1*), senescence (*TNF, CDKN1A, CDKN2A, MAPK14, IL1A, IL1B, HMGB1, MMP1, MMP12, MMP13, TIMP1, NFKB1* and *IFNG*) and T cell exhaustion (*TNFRSF18/GITR, TCF7, HAVCR2, TIGIT, LAG3*) were found to be upregulated in tumors with high PLK1 levels, potentially contributing to metastasis and therapy resistance [Fig6D]. To further corroborate our claim, we used the METABRIC study (Curtis et al., 2012; Pereira et al., 2016) to classify tumors into luminal, basal and Her2+. We found the highest expression of *PLK1* in basal tumors followed by Her2+ tumors and a similar trend was observed with PD-L1 (*CD274*) expression [FigS5B]. We have previously shown that Plk1 plays an important role in delaying tumor development as a consequence of cell and non-cell-autonomous effects. Additionally, the results displayed here highlight mechanisms that shape the phenotype of immune cells in a high CIN microenvironment to make these tumors susceptible to immune checkpoint inhibitors in a non-cell-autonomous manner. Our results are in line with previous observations that show how non-cell-autonomous effects of CIN promote tumor growth by inducing an immunosuppressive microenvironment, altering tumor antigens/defects in antigen presentation, mediating chronic inflammation and facilitating cancer immunoediting (Davoli et al., 2017; Duijf & Benezra, 2013). Future experiments aimed at using immune checkpoint inhibitors for the treatment of high CIN tumors need to be explored.

**Figure 6:**
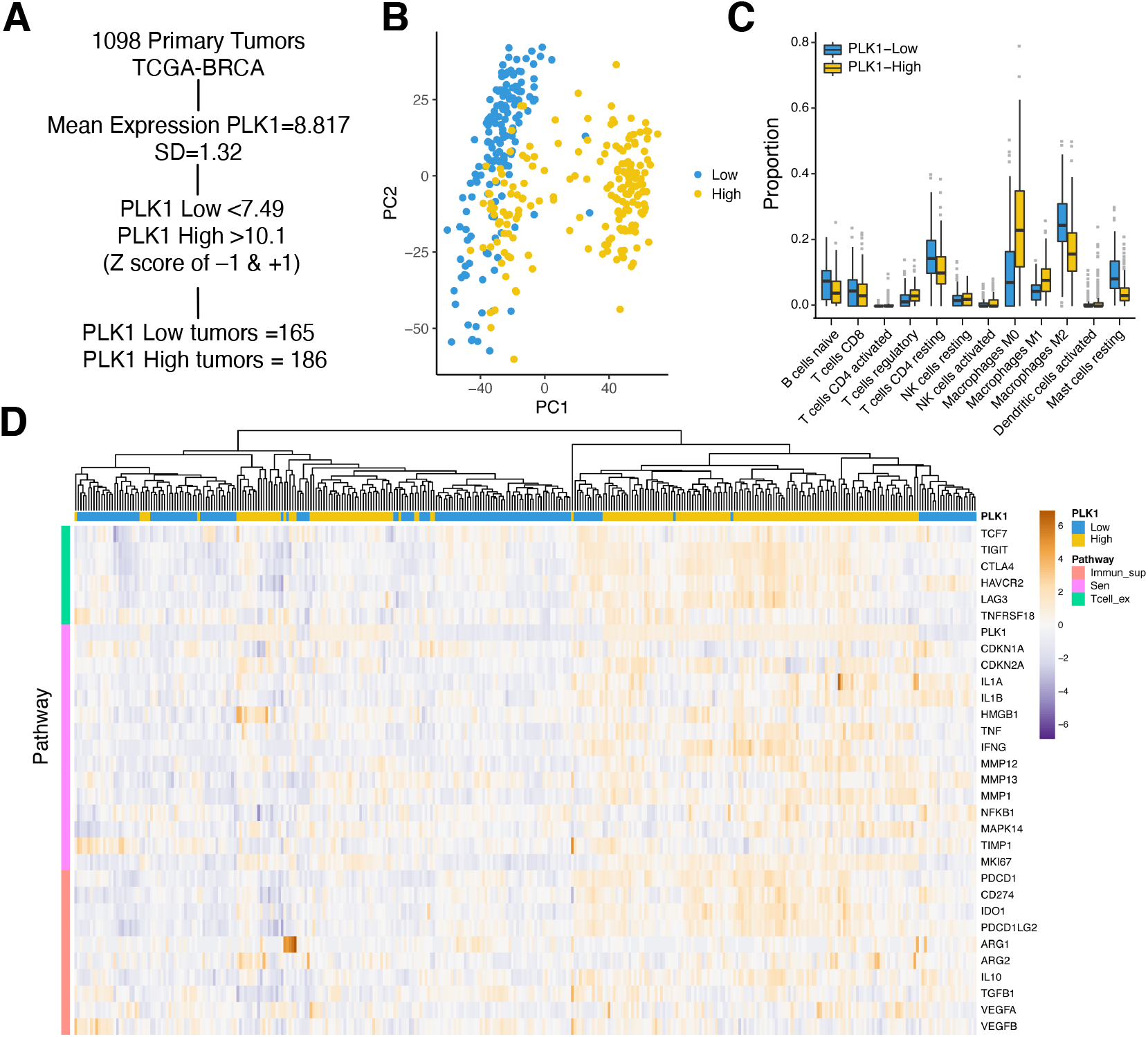
Human breast tumors of the TCGA cohort expressing high levels of PLK1 have an immunosuppressive phenotype. **A)** Schematic description of sorting PLK1 high versus PLK1 low tumors from the TCGA-BRCA cohort. A total of 186 (PLK1 high) and 165 (PLK1 low) tumors with a Z-score of +1 and –1, respectively, were identified. **B)** Unsupervised clustering of PLK1 high versus PLK1 low tumors using principal component analysis. Samples color-coded based on the groups (PLK1 low: blue, PLK1 high: yellow). **C)** Cellular deconvolution of PLK1 high and PLK1 low tumors using relative CIBERSORT to characterize the heterogeneity in immune cell composition of tumors with different levels of CIN. **D)** Heatmap showing gene expression data (Log_2_norm_count+1) and clustered into three different signatures associated with senescence, T-cell exhaustion and immune suppression from both PLK1 high (yellow) and PLK1 low (blue) tumor cohorts.

## Discussion

CIN is a hallmark of cancer known to drive immune evasion (Davoli et al., 2017). A crucial question in the field is how different levels of CIN influence this phenomenon, as this information could help in identifying effective adjuvant therapies for various cancers.

In this study, we show that overexpression of Plk1 in the mammary gland induces CIN. This results in a SASP phenotype, with aneuploid senescent tumor cells leading to a chronic inflammatory, pro-tumorigenic and aged microenvironment (Andriani et al., 2016; Q. He et al., 2018; Rao et al., 2017). We found that Plk1 tumors showed an immune evasive phenotype with enhanced RELB/p38a signaling, upregulation of PD-L1, decrease in cytotoxic NK cells and increased infiltration of Tregs. Lastly, we also revealed that PLK1 high breast tumors showed significantly increased expression of immune checkpoint and T cell exhaustion markers. This could result in a better response to immune checkpoint inhibitors, although these findings from computational studies need further validation. CIN can promote immune detection (Watson & Elledge, 2017) and based on the tumor milieu of immune cells and their phenotype at the early stages we show how high CIN modulates the tumor microenvironment to transition to an immune evasive state.

Aneuploidy triggers an innate immune response (Santaguida et al., 2017) and NK cells mediate the clearance of cells with abnormal karyotypes by activating NKG2D receptors or responding to a SASP gene expression signature (Raulet & Guerra, 2009). Validation of a similar phenomenon by blocking NK cells in vivo showed reduction in survival time of aneuploid tumors, suggesting that NK cells not only mediate immune sensing, but could also favor a suppressive microenvironment. Future experiments showing the interaction between NK cells and other immune cells at different stages of tumor progression would explain the comprehensive (dual) role of NK cells in aneuploid sensing. However, in an immunosuppressive microenvironment, NK cell infiltration is reduced (Davoli et al., 2017). Additionally, NK cells become dysfunctional through the exposure of inhibitory receptors (Cózar et al., 2021). Plk1 expression during early stages of tumor development drives NK cells to a mature state (CD11b^−^CD27^+^) with reduced effector capabilities compared to tumors with low CIN, although the activation of NKG2D showed no significant differences. We identified *Socs3* (Meisam et al., 2018), *Irf8* and *Cx3cr1* (Mace et al., 2016; Ponzetta et al.,2013) as key molecules affecting the maturation, proliferation, cytotoxic function and infiltration of these cells. Moreover, Plk1 overexpressing tumors activate both the canonical and non-canonical NF-kB pathway. Thus, our findings support the previous notion which underlines the importance of NF-kB signaling in chromosomally unstable cells for the recruitment of NK cells to eliminate them (Wang et al., 2021). Since high levels of aneuploidy result in cell cycle arrest and senescence which recruits NK cells, and since SASP (anti- and pro-tumorigenic) is under tight regulation of NF-kB, the components of SASP could essentially drive the functional capabilities of NK cells *in vivo*.

Macrophages are an important innate immune cell type that dominates the aneuploid tumor microenvironment. They act as a double-edged sword, either by facilitating phagocytosis of tumor cells (M1) or by inducing angiogenesis and promoting tumor invasion (M2). The balance between these two may impact tumor progression and therapy outcome (Lewis & Pollard, 2006). Chromosome 9p copy number gains involving PD-L1 upregulate PLK1 across major cancer types (Budczies et al., 2017) and PD-L1 expression in tumors is often associated with infiltration of immune suppressive macrophages and neutrophils (Zhu et al., 2020). Similarly, PD-L1 expression correlates with an EMT phenotype and is transcriptionally regulated by non-canonical NF-kB signaling (Asgarova et al., 2018) with TNFα and TGFβ as key drivers (Thiery & Sleeman, 2006). Interestingly, our study shows an enrichment of Arg1^+^ TAMs subset in Her2-Plk1 tumors with upregulation of gene sets involved in TGFβ signaling, reactive oxygen (ROS) and EMT signatures, which drive early modulation of immune responses. It has to be noted that Her2 tumors are also aneuploid, although to a lesser degree and Arg1^+^ TAMs were not observed during early stages of tumor development. Simultaneous targeting of Arg1 and other associated proteins involved in EMT may provide a promising opportunity to repolarize macrophages, thereby increasing the efficacy of checkpoint blockade in high CIN tumors. We predict SASP to act as an upstream mediator regulating the effector abilities of NK cells, polarizing TAMs to contribute to an EMT phenotype inducing the expansion of FoxP3^+^ Tregs in high CIN tumors, with the resulting tumor microenvironment upregulating PD-L1 expression at later stages of tumor progression.

Cytokine production is a key event in immune response during stress induced inflammation and correlates negatively in autoimmune disorders and cancer. PLK1 is known to contribute to the pathogenesis of SLE (Systemic lupus erythematosus) by activation of Aurora-A/PLK1/mTOR signaling pathway and autoantibody production, besides PLK1 itself being upregulated in splenic B220^+^ cells of lupus mice (Y. Li et al., 2022). Our study underscores the importance of several identified markers associated with altered metabolism and dysregulated signaling in B cells in high CIN tumors. Future studies involving identification of clinical manifestations and blockade of B-cell responses in PLK1 overexpressing tumors ought to be explored as a potential target.

We found that different innate and adaptive immune cells contribute to immune suppression in the context of high CIN tumors that express PLK1. Non-cell-autonomous effects such as induction of SASP in high CIN tumors could be mediated by non-canonical NF-kB (RELB) signaling. Although we have previously reported a good prognosis associated with high levels of PLK1 we further show that non-cell-autonomous effects mediated by SASP modulate the immune cells perhaps making high CIN tumors refractory to existing treatments at later developmental stages, and adding PLK1 as a potential oncogene in cancer studies. Understanding the immune repertoire of tumors expressing high levels of CIN will aid in identifying better therapeutic strategies involving immune checkpoint or NF-kB inhibitors in combination with existing first-line treatments.

## Supplementary Figure Legends

**FigS1: Bulk genomic and transcriptomic analysis of Her2 and Her2-Plk1 tumors. A)** Relative number of structural variants per individual chromosome in mammary tumors of Her2 and Her2/Plk1 groups. **B)** Percentage of tumor cells in Her2 and Her2-Plk1 with the indicated mitotic errors. **C)** Heatmap visualization of hierarchial clustering in Her2 and Her2-Plk1 tumor samples (n=3). **D)** GO functional analysis of differentially expressed genes retrieved using DAVID. The top 20 significantly (P<0.05) enriched GO terms in biological processes are represented. The size of the dot is based on the gene count enriched in the pathway and the color of the dot represents the pathway adjusted enrichment score.

**FigS2: Non-cell-autonomous effects of Plk1. A)** Immunodetection of proteins from lysates of rtTA stable MCF7 cells (estrogen receptor expressing) and **B)** Cal51 cells (Triple negative) were generated with and without PLK1, where transcription of PLK1 is regulated by doxycycline. The cells were treated with doxycycline for 24, 48 and 72hrs. Protein lysates from both cell lines were immunoblotted with anti-p38, anti-NF-kB-p65, anti-RELB with anti-GAPDH as a loading control. **C**) Expression of P38a and NF-kB-p65 in breast tumors from Her2 and Her2-Mad2 mice. Tumor cell lysates were used for immunodetection by using anti-NFkB-p65, anti-RELB, anti-P38, anti-pP38 and actin as a loading control.

**FigS3: Chromosome copy numbers in Her2 and Her2-Plk1 samples detected by FISH and comparison of single-cell RNA sequencing between both groups. A)** Hematoxylin and Eosin staining of early-stage mammary glands in both Her2 (16 days on doxycycline) and Her2-Plk1 (22 days on doxycycline) mice. **B)** Hematoxylin and Eosin, PLK1 and fluorescence in situ hybridization (FISH) using two centromeric probes against chromosomes 16 and 17 on paraffin sections of early-stage mammary glands. P=0,0073, unpaired t-test. Scale bar: 50μm. **C)** Graphs displaying the total number of genes, total number of unique molecular identifiers (UMIs) and percentage of mitochondrial RNA per cell for the Her2 and Her2-Plk1 samples from single-cell RNA-sequencing experiments after filtering for cells carrying the information from more than 200 genes and mitochondrial content of less than 10% (upper panels). Similarly, distribution of the number of detected genes, number of unique molecular identifiers and percentage of mitochondrial RNA combined for all samples displayed by dimensionality reduction by UMAP (lower panels). **D)** Immune cell populations by sample type (Her2: blue, Her2-Plk1: yellow) are shown by dimensionality reduction. **E)** Relative numbers of different immune cell populations across Her2 (blue) and Her2-Plk1 (yellow) overexpressing tumors.

**FigS4: Cell type annotations in single-cell RNA-sequencing experiments and gene expression in two types of macrophage subsets. A)** Heatmap showing normalized gene expression levels of cluster-specific marker genes detected via differential expression between the clusters. **B-C)** Violin graphs showing the normalized gene expression levels of TNF family related inflammatory genes in both CD11b^+^CD11c^−^ (B) and CD11b^+^CD24^−^ (C) macrophage subsets.

**FigS5: Copy number changes of the human breast tumors of the TCGA cohort and gene expression levels in the METABRIC cohort. A)** Violin graphs showing the percentage of copy number gain or loss in low PLK1 breast tumors and high PLK1 tumors from the TCGA cohort. Each arm of the chromosome is plotted and represented by one dot. *, p<0.05; **, p<0.005, One way ANOVA, Tukey’s multiple comparison test. **B**) Gene expression levels of *PLK1* and *CD274* shown for three different subtypes of tumors, luminal (1865 samples), basal (335 samples), and HER2+ (247 samples), Welch t-test.

## Methods

### Mouse models

*TetO-rat-Her2*, and *ColA1-Plk1* heterozygous animals were bred to *MMTV-rtTA* (Carcer et al., 2018), and only heterozygous female animals for all transgenes were used in this study. Breeding and experimentation were performed at the DKFZ animal facilities, with ethical approval from the corresponding Animal Welfare and ethical review bodies under the protocol number G231/15 until December 2020 and subsequently with protocol number G-18/21. Eight-week-old female mice were fed with 625ppm doxycycline-enriched diet to exclusively express the transgenes in the mammary gland.

For genotyping the mice, isolation of tail-DNA was performed by incubation in 0.05 M NaOH at 98 °C for 1.5 h and subsequent neutralization with 1 M Tris HCl pH 7.5. The following oligonucleotides were used to genotype the ColA1-Plk1 allele: KH2-Plk1 A: 5′-GCACAGCATTGCGGACATGC-3′, KH2-Plk1 B: 5′-CCCTCCATGTGTGACCAAGG-3′, KH2-Plk1 C: 5′-GCAGAAGCGCGGCCGTCTGG-3′. For all transgenes, the following PCR program was applied: 94 °C for 2 min, 30 times [95 °C for 30 s, 60 °C for 30 s, 72 °C for 30 s], and a final step at 72 °C for 1 min.

8 week-old female mice were treated with an NK blocking antibody (500µg of NK1.1 blocking AB on day 1, followed by 250µg of the same antibody on day2 to remove NK cells. Mice were then kept on doxycycline and given 250µg of NK1.1 blocking antibody every week to maintain the depletion state, until tumors were observed.

### Bulk RNA Sequencing

Total RNA was isolated from tumor tissues using buffer RLT from the RNeasy Mini kit (Qiagen) and RNeasy spin columns (Qiagen, Venlo, The Netherlands) according to the manufacturer’s protocols. Libraries were prepared using the Illumina TruSeq Stranded mRNA Library Prep (Illumina, San Diego, California, USA) according to the manufacturer’s protocols. The libraries were sequenced as 50 base pair (bp) single-end reads using HiSeq 4000 (Illumina). The sequenced reads were aligned to the mm10 reference genome using kallisto v0.46.1 (Bray et al., 2016). Raw counts were normalized and differentially expressed genes (DEGs) were calculated in R by using the DEseq2 package (Love et al., 2014). GO analysis and GSEA of differentially expressed genes was performed using the packages ‘ClusterProfiler’ (Wu et al., 2021), ‘enrichplot’(G, 2021) and ‘pathview’(Luo & Brouwer, 2013) in R, with P<0.05 being a statistically significant difference.

### Cytokine Array

Cytokine levels were analyzed by using the Proteome Profiler Mouse Cytokine Array Panel A kit (R&D Systems, Minneapolis, MN, USA), following the manufacturer’s instructions. In brief, tissues were lysed using lysis buffer (containing protease inhibitor cocktail). Protein concentration in the lysate was measured using a BCA protein assay kit (Thermo Scientific, Waltham, MA, USA). Tissue protein diluted in array buffer was incubated with the ready-to-use pre-coated array membranes overnight at 4°C on a rocking platform shaker. The membrane was washed and incubated with streptavidin–horseradish peroxidase (HRP) buffer for 30min. Next, the membrane was washed and incubated with the Chemi Reagent mixture at 23–27°C for 1min and analyzed using the LAS-300 system (Fujifilm, Tokyo, Japan). Dot density was analyzed using Fiji 3.0.

### ß-galactosidase Activity Assay

Senescence in the tumor tissues was determined by using the ß-Galactosidase (ß-Gal) Activity Assay Kit (Fluorometric) from Biovision (K821-100). Tissues were homogenized in assay buffer following the manufacturer’s instructions and a standard curve was plotted with the tumor samples and standard solution (supplied in the kit). Fluorescence was measured (Ex/Em= 480/530) in kinetic mode, choosing two time points (T1 & T2) in the linear range to calculate the ß-gal activity. Quantified ß-gal activity is expressed in the unit µU/mg of protein.

### Flowcytometry and Sorting

Her2 and Her2-Plk1 tumors were digested using 100 U/ml collagenase IV and 50 u/ml DNase I in complete RPMI medium at 37°C. To determine the percentages of immune cell subsets, single-cell suspensions from spleen and tumor tissues were incubated with anti-CD45, anti-Cd3, anti-Ly6C, anti-CD4, anti-CD8, anti-CD11c, anti-MHCII, anti-CD11b, anti-Nk1.1, anti-Nkp46 and anti-CD25 (all from Biolegend) and anti-Foxp3 (eBioscience). For quantification of intracellular Foxp3, cells were incubated with permeabilization buffer (Foxp3 transcription factor staining buffer kit, eBioscience) for 4 to 5 hours and then washed, fixed, and permeabilized with perm/fix buffer (eBioscience). For the analysis, events were collected on a BD FACS Canto (Becton, Dickinson& Company) cytometer and all results were analyzed with FlowJo (Version10.1) software. For sorting of mammary glands at early stages, fresh tissues were minced and incubated in complete RPMI 1640 medium in the presence of collagenase IV (100 U/ml) and DNase I (50 U/ml). They were incubated in gentleMACS Octo Cell Disassociator (Miltenyi Biotec) at 37°C with gentle agitation. Medium was then added to wash the cells and neutralize the reaction. Samples were filtered twice, stained with anti-CD45 and anti-EpCAM antibodies for subsequent isolation of hematopoietic cells and mammary epithelial cells. The cells were sorted on a FACS Aria II (Becton, Dickinson& Company) cell sorter using the gating strategy illustrated in Fig. 3A.

### Immunohistochemistry

Immunohistochemistry on mouse tissues was performed on formalin-fixed paraffin-embedded sections. Following deparaffinization with xylene and rehydration with graded ethanol, antigen retrieval was performed using 0.09% (v/v) unmasking solution (Vector Labs) for 30 min in a steamer. Inactivation of endogenous peroxidases was carried out using 3% hydrogen peroxide (Sigma) for 10 min. Secondary antibody staining and biotin-streptavidin incubation were performed using species-specific VECTASTAIN Elite ABC kits (Vector Labs). DAB Peroxidase Substrate kit (Vector Labs, SK-4100) was utilized for antibody detection. Primary antibodies used were anti-Plk1 (1:20), anti-PD-L1 (1:150) and anti-CD206 (1:500). Sections were then subjected to sequential incubations with the indicated biotinylated secondary antibodies and avidin/biotinylated enzyme reagents (ABC kit (PK-6101; PK-61-04)). For PD-L1 and CD206, the quantitation was performed using the StrataQuest software (TissueGnostics) to determine the total sum area of positive cells.

### Immunodetection

For protein extraction of mouse tissues, MCF7-rtTA, MCF7-rtTA-Plk1, Cal51-rtTA and Cal51-rtTA-Plk1 cells were lysed in Laemmli lysis buffer (0.25 M Tris-HCl pH 6.8, 2.5% glycerol, 1% SDS, and 50 mM DTT). Samples were then boiled for 10 min and cleared by centrifugation. Proteins were separated on 12% Tris-Glycine acrylamide gels (BioRad), transferred to nitrocellulose membranes (BioRad), and probed using the following specific antibodies: anti-PLK1(1:500), anti-NF-kB-p65 (1:1000), anti-RELB (1:1000), anti-p38a (1:1000), anti-pp38MAPK T180/Y182 (1:1000), anti-vinculin (1:2000) and anti-actin (1:2000). After incubation with anti-mouse/anti-rabbit secondary antibodies, respectively, the signal was detected by using SuperSignal chemiluminescence substrate (Thermo-scientific) and the iBright1500 system (Invitrogen).

### Fluorescence In Situ Hybridization (FISH)

FISH was performed on formalin-fixed paraffin-embedded 5-µm sections after deparaffinization, rehydration, and antigen retrieval with unmasking solution 0.09% (*v/v*) (Vector Labs, California, USA, NA). The BAC DNAs were labeled by nick translation according to standard procedures. A probe mix was prepared with 3 µL of the labeled probe and 7 µL Vysis LSI/WCP Hybridization buffer. Hybridization was performed using the Abbott Molecular Thermobrite system with the following program: Denaturation at 76 °C for 5 min, hybridization at 37 °C for 20–24 h. Pan-centromeric probes were made using pairs of BAC clones for each chromosome: Chr 16-RP23–290E4 and RP23– 356A24 labeled with SpectrumOrange-dUTP (Vysis), and Chr 17-RP23–354J18 and RP23– 202G20 labeled with SpectrumGreen-dUTP (Vysis). A total of at least 400 cells from each sample with an average of 10 ROIs from each sample was considered for quantification of levels of aneuploidy in both, Her2 and Her2-Plk1, samples.

### Tumor cell culture

Mammary tumors at endpoint were digested with the Mouse Tumor Dissociation Kit (130-096-730, Miltenyi). The gentleMACS Dissociator (130-095-937, Miltenyi) was used to disgregate the tissues. For time-lapse microscopy, cells were cultured on 8 well chambered coverglass (Thermo Scientific, 155411) in DMEM supplemented with 10% tet-Free Serum, 4mM L-glutamine (GIBCO), 100µg/ml Penicillin/Streptomycin (GIBCO), 5µg/ml insulin (Sigma), 10µg/ml EGF (Sigma), 1µg/ml hydrocortisone (Sigma), 1µM progesterone (Sigma), 5µg/ml prolactin (NIHPP), and 1µg/ml doxycycline. Imaging was performed on a Zeiss Cell Observer with 2μm optical sectioning across 18μm stack, 30 frames/h. Zeiss Zen 2 software served for image analysis.

### Human Cell Line Culture

MCF7 cells were cultured in DMEM with 10% FBS (Gibco) and 1% penicillin/streptomycin (Life Technologies), non-essential amino acids (0.1 mM, Life Technologies), insulin (10 ug/mL) and sodium pyruvate (1 mM, Life Technologies). Cal51 cells were maintained in DMEM high glucose supplemented with 10% FBS (Gibco), 1% Penicillin-Streptomycin (Life Technologies) and 1% glutamine (Life Technologies). To generate PLK1-inducible cell lines, MCF7 were first infected with a rtTA-expressing retrovirus and selected with puromycin (1 µg/ml) whereas Cal51 cells expressing rtTA were sorted for GFP positive cells after infection. These cells were then infected with an inducible Tet-ON lentivirus carrying the human PLK1 cDNA (pLenti CMVtight Hygro DEST from Addgene #26433) and selected with hygromycin (350 µg/ml). The cells were incubated with and without doxycycline (5mg/ml) for 48 h to detect the expression of p38 and NF-kB related genes.

### DNA Sequencing and Analysis

Snap frozen tissue was used for extraction of genomic DNA using the DNeasy Blood & Tissue Kit (Qiagen) as per manufacturer guidelines. Library preparation and low-coverage sequencing were performed on an Illumina HiSeq 2500 platform (Illumina) using 50-base pair single-end reads. The DKFZ Genomics and Proteomics core facility handled library preparation and sequencing. For the analysis of the sequencing data, the reads were aligned to the mm 10 build of mouse reference genome using a Burrows-Wheeler Aligner (BWA; version 0.7.10). Tumor coverage files were log2-normalized to mouse genomic DNA derived from normal mammary tissue. Circular binary segmentation (CBS; R package) was applied, and somatic copy number alterations were manually characterized as detailed: Whole chromosome gains/losses were defined as chromosome-wide shifts in the segmentation of a chromosome, whereas partial chromosome gains/losses entailed changes spanning at least one-fifth of the chromosome. Focal amplifications and deletion encompassed events smaller than this. When the number of copy number state switches on a chromosome exceeded ten, they were called gross chromosomal rearrangements.

### TME Cell Type Inference

The relative fractions of 22 cell subsets within the tumor microenvironment (TME) lymphocyte population were inferred from batch-normalized gene-expression profiles for each tumor sample using CIBERSORT v.1.06 [https://cibersort.stanford.edu/] (Newman et al., 2019). The program was run with the default LM22 reference matrix to infer the relative number of immune cells in samples expressing high and low levels of PLK1.

### Human transcriptomic data sets and pre-processing

Gene expression data of PLK1 for 1108 breast cancer samples from TCGA was obtained in the form of Log_2_ normalized counts+1 from the Xena browser (Goldman et al., 2020). It was used to stratify cohorts of tumors as PLK1 high and PLK1 low based on the Z-score of the expression data. The corresponding 165 tumor samples (PLK1 low) and 186 tumor samples (PLK1 high) were downloaded from GDC’s legacy archive, normalized, and filtered using the R/Bioconductor package TCGAbiolinks version 2.22.3 using GDCquery, GDCdownload, and GDCprepare functions for tumor types (level 3, and platform “IlluminaHiSeq_RNASeqV2”). Unsupervised clustering by PCA was done to identify the relation between the groups.

PLK1 and PD-L1 expression was additionally analyzed on a larger dataset, the METABRIC cohort (Curtis et al., 2012; Pereira et al., 2016). We stratified the tumors based on their ER and HER2 status and compared 1865 luminal, 335 basal, and 247 HER2+ samples, Microarray z-scores were used for gene expression level comparison.

### Single-cell RNA sequencing

Mammary glands of mice 16 and 22 days after doxycycline administration were isolated and dissociated into single-cell suspensions in the presence of collagenase IV (100 U/ml) and DNase I (50 U/ml). The suspension was incubated in a gentleMACS Octo Cell Disassociator (Miltenyi Biotec) at 37°C with gentle agitation. Leukocytes were selected by staining with an anti-mouse CD45 antibody and collected with the help of a sorter (FACS Aria II). Cells were then subjected to single-cell RNA sequencing by using the 10x Genomics platform. Reverse transcription and library preparation were performed by following the protocol Chromium Next GEM sc 3 Reagent v3.1, followed by paired-end sequencing runs on the platform Illumina NovaSeq 6000 S1. Sequencing data were processed by using the 10x Genomics Cell Ranger 3.0.0 (Zheng et al., 2017). The data were further analyzed by using the R package Seurat (Butler et al., 2018). Cells were selected for further analysis if they carried more than 200 genes and showed a mitochondrial content of less than 10%. By principal component analysis and manual inspection of the corresponding ‘Elbow graph’, the first 30 features were selected for downstream clustering and dimensionality reduction. Clusters were determined by a shared nearest neighbor modularity optimization based on the Louvain algorithm (Waltman & Eck, 2013)with a resolution of 0.5. The SingleR package was used to annotate cells to a reference database (ImmGen) (Consortium et al., 2008). Differential gene expression between clusters and between samples within individual clusters was analyzed by using a Wilcoxon Rank Sum test. Gene set enrichment analysis was performed by using the R package clusterProfiler (Yu et al., 2012) with the datasets for hallmark genes (H), curated genes (C2), gene ontology genes (C5), and immunologic signatures (C7). The data were visualized by using the R package ggplot2 (Wickham, n.d.) and the Python packages matplotlib and seaborn (Waskom, 2021).

### Statistical analysis

GraphPad Prism 8 was used for statistical testing, excluding the single cell analysis.

### Data availability

The data that support the findings of this study are available within this article and Supplementary Files, or available from the authors upon request.

## Acknowledgments

We would like to thank members of the Sotillo lab for their valuable inputs during the discussion and writing of the manuscript. We would further like to thank M. Simon, N. Beumer, and S. Kotrbaty for insightful discussions about data analysis and G.de Carcer and G. Cui for their helpful suggestions. We thank also the DKFZ Core Facilities of Light Microscopy, Flow Cytometry, and Genomics and Proteomics for the excellent technical assistance; and the Central Animal Laboratory for animal husbandry. We appreciate the help of Jan-Philipp Mallm and the DKFZ Single-Cell Sequencing Open Lab in designing and conducting the scRNA-Seq experiments. Schemes have been generated with BioRender.com.

## Author Contributions

S.K. and R.S. designed the study and corresponding experiments. S.K. performed the experiments and analyzed data. L.V. developed code and performed the analysis of the single-cell RNA-sequencing data. J.K. provided code for the analysis of single-cell data and B.B. and C.I. supervised it. M.R. performed in vitro experiments and S.C. performed and analyzed FISH. S.K., L.V., and R.S. wrote the manuscript. R.S. acquired funding and performed supervision. All authors read and contributed to the manuscript.

---

**Supplementary Table 1.** Number of cells analyzed per sample and cell type.

| Number of cells | *Her2* | *Her2-Plk1* | Total |
| --- | --- | --- | --- |
| Follicular B cells | 2062 | 605 | 2667 |
| CD4+ T cells | 7808 | 2312 | 10120 |
| Regulatory T cells | 608 | 225 | 833 |
| CD8+ T cells | 4519 | 1043 | 5562 |
| CD8+ effector T cells | 239 | 73 | 312 |
| CD11b+24- macrophages | 120 | 15 | 135 |
| CD11b+11c- macrophages | 1082 | 71 | 1153 |
| Macrophages steady-state | 468 | 67 | 535 |
| Natural killer cells (CD3-, NK1.1+) | 950 | 118 | 1068 |
| CD8+ plasmacytoid dendritic cells | 121 | 36 | 157 |
| Double negative dendritic cells | 34 | 9 | 43 |
| Neutrophils (arthritic) | 349 | 30 | 379 |
| Mast cells (MC.TR) | 106 | 22 | 128 |
| Fibroblasts | 24 | 4 | 28 |
| **Total** | **18490** | **4630** | **23120** |

\*Shown is the number of cells after filtering of cells for mitochondrial content and number of detected genes.

**Supplementary Table 2.** Total number of luminal and basal tumors in PLK1 high and PLK1 low tumors from TCGA.

| **Tumor Type – PLK1 High** | ***Total number*** |
| --- | --- |
| Breast invasive ductal carcinoma (basal tumors) | 171 |
| Breast invasive lobular carcinoma (luminal cancers) | 7 |
| Basal cell carcinoma | 1 |
| Metaplastic breast cancer | 3 |
| Breast invasive ductal + luminal carcinoma | 2 |
| **Tumor Type- PLK1 Low** | ***Total number*** |
| Breast invasive ductal carcinoma (basal tumors) | 68 |
| Breast invasive lobular carcinoma (luminal cancers) | 84 |
| Basal cell carcinoma | - |
| Metaplastic breast cancer | - |
| Breast invasive ductal + luminal carcinoma | 9 |

